# Prognostic biomarkers in oral squamous cell carcinoma: a systematic review

**DOI:** 10.1101/164111

**Authors:** César Rivera, Ana Karina de Oliveira, Rute Alves Pereira e Costa, Tatiane De Rossi, Adriana Franco Paes Leme

## Abstract

Over the years, several tumor biomarkers have been suggested to foresee the prognosis of oral squamous cell carcinoma (OSCC) patients. Here, we present a systematic review to identify, evaluate and summarize the evidence for OSCC reported markers. Eligible studies were identified through a literature search of MEDLINE/PubMed until January 2016. We included primary articles reporting overall survival, disease-free survival and cause-specific survival as outcomes. Our findings were analysed using REporting recommendations for tumor MARKer prognostic studies (REMARK), QuickGo tool and SciCurve trends. We found 41 biomarkers, mostly proteins evaluated by immunohistochemistry. The selected studies are of good quality, although, any study referred to a sample size determination. Considering the lack of follow-up studies, the molecules are still potential biomarkers. Further research is required to validate these biomarkers in well-designed clinical cohort-based studies.

## INTRODUCTION

Oral squamous cell carcinoma (OSCC) is the most common malignancy of the head and neck (excluding nonmelanoma skin cancer), with more than 300,000 new cases reported annually worldwide[1]. The disease has a high morbidity rate (37.8%) five years after diagnosis (http://www.cancer.gov/statistics/find-2003-2009data); despite the progress in research and therapy, survival has not improved significantly in the last few decades [2]. The search for prognostic markers represents a continuing challenge for biomedical science.

A cancer biomarker may be a molecule secreted by a tumor cell or a specific response of the body to the presence of cancer [3]. Biomarkers can be used for patient assessment in multiple clinical settings, including estimating the risk of disease and distinguishing benign from malignant tissues [4]. Cancer biomarkers can be classified based on the disease state, including predictive, diagnosis and prognosis biomarkers [5]. A prognostic biomarker informs about a likely cancer outcome (e.g., overall survival, disease-free survival, and cause-specific survival) independent of treatment received [6].

According to the NCI Dictionary of Cancer Terms (https://www.cancer.gov/publications/dictionaries/cancer-terms) the overall survival (OS)corresponds to the length of time from either the date of diagnosis or the start of treatment for cancer, which patients diagnosed with the disease are still alive. Disease-free survival (DFS, also called relapse-free survival) offers the length of time after primary treatment ends that the patient survives without any signs or symptoms of that cancer. Cause-specific survival (CSS) is the length of time from either the date of diagnosis or the start of treatment for cancer to the date of death from the disease.

From the identification of a promising biomarker to its clinical use, there is a long pathway involving many complicated hurdles, such as estimating the number of patients needed for the validation phase and statistical validation, among others [7, 8]. This validation and qualification are responsible for linking the promising biomarker with a biological process to clinical endpoints [9].

Considering several tumor biomarkers have been suggested to predict the prognosis of OSCC patients, we performed a systematic review, which is widely accepted as a “gold standard” in medicine based on evidence [10], to identify, evaluate and summarize the evidence for OSCC reported markers.

## MATERIALS AND METHODS

We performed a systematic review to conduct this investigation. The independent variables were prognostic biomarkers; the dependent variables were OSCC outcomes.

### Search strategy

A systematic review allows critical analysis of multiple research studies. Aiming to answer the question “what are the biomarkers of OSCC?”, a systematic literature search based on keywords was performed. As PubMed comprises more than 26 million citations from the biomedical literature from MEDLINE, it is the search engine of choice to initiate queries in the health sciences. To identify all the primary research studies that evaluated candidate biomarkers in OSCC, we searched the MEDLINE/PubMed (http://www.ncbi.nlm.nih.gov/pubmed) medical literature database up to January 18, 2016. The search strategy was based on combinations of the following keywords: “mouth neoplasms" [MeSH] and "biomarkers" [MeSH] and (risk ratio [Title/Abstract] or relative risk [Title/Abstract] or odds ratio [Title/Abstract] or risk [Title/Abstract]) and ("humans"[MeSH Terms] and English [lang]).

### Inclusion criteria

Articles were included based on a previously published protocol [11]. Briefly, studies were selected if they examined the impact of a potential biological marker on at least one of the features in OSCC patients: OS, DFS or CSS. These definitions were assessed among the selected papers. In addition, if a study was focused on isolated or combined (multiple) tumor biomarkers, it must have been subjected to multivariable analysis with one or more additional variables.

### Exclusion criteria

Articles were excluded from the present review for the following reasons: i) lack of the terms “oral cancer” and “risk” in their titles, abstracts or keywords; ii) absence of risk ratios and iii) unclear defining criteria for groups and variables.

### Potential prognostic biomarker

To determine whether a biomarker is potentially prognostic, the selected articles showed: i) a formal test (binary logistic regression or Cox proportional hazards model) and ii) a statistically significantly association between the biomarker and outcome [6]. The computed risk (odds ratio,OR or hazard ratio, HR) was reported as the risk of a specific outcome from the biomarker group versus the reference group, with OR/HR>1 indicating increased risk and OR/HR<1 indicating decreased risk.

### Data extraction

One investigator reviewed all the eligible studies and carefully extracted the study characteristics, including the article citation information, biomarker name and classification, condition or outcome, laboratory technique, sample size, number of clinical outcomes, status of biomarker expression, statistical test method, computed risk and its p-value and 95% confidence interval (CI). The main biological processes in which the biomarkers are involved were obtained using QuickGo (http://www.ebi.ac.uk/QuickGO).

### Quality assessment

Quality assessment was performed in duplicate for each eligible study by three independent reviewers using operationalized prognostic biomarker reporting the REMARK guidelines [12] and extracted details on 20 items. The inter-observer agreement was evaluated using Kappa statistics.

### Publication trends

To observe the publication trends in the selected potential OSCC biomarkers, we searched the scholarly literature in SciCurve Open (http://www.scicurve.com). SciCurve Open is a search engine that transforms a systematic literature review into an interactive and comprehensible environment [13].

## RESULTS

### Studies searching for OSCC biomarkers:proteins are the most analysed molecules

The keyword search strategy identified 403 suitable abstracts, from which 320 were excluded by reviewing the title and abstract during the screen because they did not meet the eligibility criteria. Full text articles were obtained for 83 studies (34 with single markers and 49 with multiple or combined markers).

Forty-five of these articles were excluded for different reasons, including: out of goal (3 articles), unavailability online (2 articles), lack of multivariable analysis (18 articles) and model inconsistencies (22 articles). Figure 1 shows a PRISMA diagram for this review (for details, see Supplemental file S1).

**Fig. 1.**
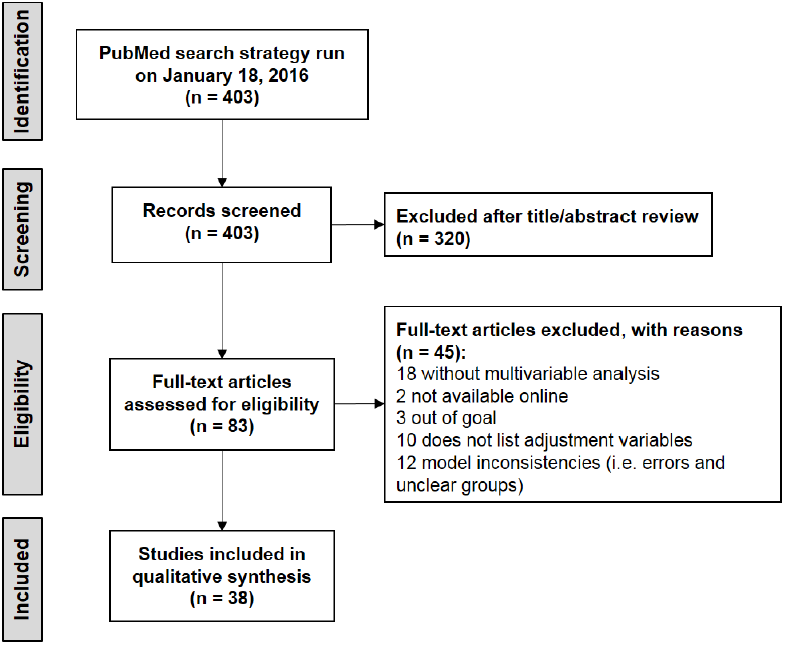
Flow diagram representing systematic literature search on biomarkers and oral cancer outcomes. Studies were included if they examined the impact of a potential biomarker on at least one of overall survival, disease free survival or cause-specific survival in oral squamous cell carcinoma patients.

The selected studies were screened, and specific study characteristics and remarks were recorded. These parameters are summarized in Table 1 (the article context is grouped according to the hallmarks of cancer [14]). Thirty-eight papers examined 41 biomarkers [15-52]. Most of them were proteins determined using immunohistochemistry (IHC) in paraffin-embedded tissues (36 of 38 studies).

**Table 1:** Characteristics of the included studies.

| Reference | Biomarker* | Change | Design and method | Study remarks |
| --- | --- | --- | --- | --- |
| **Sustaining proliferative signaling** |  |  |  |  |
| Gontarz M et al., 2014 | Proliferation marker protein Ki-67 or ki-67 (MKI67) | (+) | Poland. Retrospective. IHC. | May be useful in the selection of patients at a higher risk of recurrence who would benefit from postoperative radiotherapy |
| Ramshankar V et al. 2014 | Cyclin-dependent kinase inhibitor 2A or p16 (CDKN2A) | (+) | India. Retrospective. IHC and RT-qPCR. | CDKN2A overexpression is a single important prognostic variable in defining a high risk group. CDKN2A expression should possibly not be used as a surrogate marker for HPV infection in tongue cancers. |
|  | Human papillomavirus type 16 or HPV-16 (HPV16) | (+) |  |  |
| Tripathi SC et al., 2012 | Rho GTPase-activating protein 7 or DLC1 (DLC1) | (-) | India. Retrospective. IHC. | Loss of expression emerged as an important biomarker for predicting patients diagnosed with oral dysplasia at high risk of transformation. Is a poor prognostic marker for oral squamous cell carcinoma patients. |
| Freudlsperger C et al., 2011 | MKI67 | (+) | Germany. Retrospective. IHC. | Expression level could be used to identify a subgroup of surgically treated patients with stage I OSCC who might benefit from treatment intensification. |
| Kok SH et al., 2010 | Protein CYR61 (CYR61) | (+) | Taiwan. Retrospective. IHC. | Is a positive growth modulator of OSCC and overexpression is an independent prognostic indicator. |
| Shah NG et al. 2009 | Cellular tumor antigen p53 or p53 (TP53) | (+) | India. Retrospective. IHC. | TP53 was independently associated with DFS and OS, and CDKN2A with DFS only. |
|  | CDKN2A | (-) |  |  |
| Kim SJ et al. 2007 | Carbonic anhydrase 9 (CA9) and MKI67 combined | (+) | Korea. Retrospective. IHC. | The expression of CA9 and MKI67 may be useful for predicting prognosis in squamous cell carcinoma of the tongue. |
| Shiraki M et al. 2005 | TP53, G1/S-specific cyclin-D1 (CCND1), epidermal growth factor receptor (EGFR) combined | (+) | Japan. Retrospective. IHC. | Simultaneous expression of these markers in oral cancers might prove to be a useful indicator for identification of low- or high-risk patients. |
| Myo K et al., 2005 | CCND1 | (+) | Japan.<br>Retrospective.<br>FISH. | Aberrations in gene numbers appear to be valuable in identifying patients at high risk of late lymph node metastasis in stage I and II OSCCs. |
| Pande P et al. 2002 | TP53 | (+) | India.<br>Prospective.<br>IHC. | RB1 loss and TP53 overexpression may serve as adverse prognosticators for disease free survival of the patients. |
|  | Retinoblastoma-associated protein (RB1) | (-) |  |  |
| Bova RJ et al. 1999 | CCND1 | (+) | Australia.<br>Retrospective.<br>IHC. | CCND1overexpression and loss of CDKN2A expression predict early relapse and reduced survival in squamous cell carcinoma of the anterior tongue |
|  | CDKN2A | (-) |  |  |
| Evading growth suppressors |  |  |  |  |
| Pérez-Sayáns M et al., 2014 | Myc proto-oncogene protein (MYC) | (+) | Spain.<br>Retrospective.<br>IHC. | Its determination can be valuable when used together with other markers to assess the prognosis of OSCC patients. |
| Liu W et al. 2013 | Retinal dehydrogenase 1 or ALDH1 (ALDH1A1) | (+) | China.<br>Prospective.<br>IHC. | Expression of cancer stem cell markers ALDH1A1 and PROM1 correlate with a high risk of malignant transformation in a large series of patients with premalignant oral leukoplakia. |
|  | Prominin-1 or CD133 (PROM1) | (+) |  |  |
| Feng JQ et al. 2013 | ALDH1A1 | (+) | China.<br>Retrospective.<br>IHC. | Expression pattern was associated with malignant transformation, suggesting that it may be valuable predictors for evaluating the risk of oral cancer. |
| Suzuki F et al., 2005 | Protein S100-A2 (S100-A2) | (-) | Japan.<br>Retrospective.<br>IHC. | Patients with stage I or II invasive OSCC without expression should be considered a high-risk group for late cervical metastasis when a wait-and-see policy for the neck is being considered. |
| Tsai ST et al., 2005 | S100-A2 | (-) | Taiwan.<br>Retrospective.<br>IHC. | Loss of nuclear expression may serve as an independent prognostic marker for early-stage oral cancer patients at high risk of recurrence. A more aggressive treatment modality and intensive follow-up may be recommended for the patients with reduced expression of in tumor cell nuclei. |
| Resisting cell death |  |  |  |  |
| Moura IM et al., 2014 | Cell division cycle protein 20 homolog (CDC20) | (+) | Portugal.<br>Retrospective.<br>IHC. | High expression is associated with poor prognosis in OSCC, may be used to identify high-risk OSCC patients, and may serve as a therapeutic target. |
| Tang JY et al., 2013 | Microtubule-associated proteins 1A/1B light chain 3A or LC3 (MAP1LC3A) | (+) | Taiwan. Retrospective. IHC. | Elevated expression, which corresponds to increased level of autophagy activity, is a frequent event and an indicator of poor prognosis in human OSCC. |
| de Carvalho-Neto PB et al. 2013 | Tumor necrosis factor receptor superfamily member 6 or FAS (FAS) | (-) | Brazil. Retrospective. IHC. | DFS and CSS were significantly correlated with FAS/FASL expression profiles. The high risk category was an independent marker for earlier disease relapse and disease-specific death. |
|  | Tumor necrosis factor ligand superfamily member 6 or FASL (FASLG) | (-) |  |  |
| Inducing angiogenesis |  |  |  |  |
| Yanagawa T et al., 2004 | Heme oxygenase 1 (HMOX1) | (-) | Japan. Retrospective. IHC. | Could be used clinically as a marker for tumors possessing the potential for lymph node metastasis. This method could prove useful as an adjuvant method to detect lymph node metastasis and may help reduce the number of surgeries by indicating when surgery is unnecessary. |
| Activating invasion and metastasis |  |  |  |  |
| de Vicente JC et al., 2013 | Podoplanin (PDPN) | (+) | Spain. Retrospective. IHC. | Could be a valuable biomarker for risk assessment of malignant transformation in patients with oral leukoplakia along with histological assessment |
| de Vicente JC et al. 2012 | Src substrate cortactin (CTTN) and focal adhesion kinase 1 (PTK2) combined | (+) | Spain. Retrospective. IHC. | Strong immunoexpression of CTTN and PTK2, and not only one of them, is a predicting factor for increased cancer risk in oral premalignant lesions. |
| Hamada T et al., 2012 | Mucin-4 (MUC4) | (+) | Japan. Retrospective. IHC. | Overexpression is an independent factor for poor prognosis of patients with OSCC; therefore, patients with OSCC showing positive expression should be followed up carefully. |
| Ma LW et al., 2012 | Catenin delta-1 (CTNND1) | (+) | China. Retrospective. IHC. | May serve as a useful marker for the identification of a high risk of potentially malignant oral lesions progressing to OSCC |
| Marsh D et al. 2011 | Actin, aortic smooth muscle or SMA (ACTA2) | (+) | UK. Retrospective. IHC. | An positive, myofibroblastic stroma is the strongest predictor of OSCC mortality. |
| Zhang Z et al. 2011 | Interstitial collagenase or MMP-1 (MMP1) | (+) | China.<br>Retrospective.<br>IHC. | Up-regulation of MMP1, MMP2 might be important features of OSCC progression. |
|  | 72 kDa type IV collagenase or MMP-2 (MMP2) | (+) |  |  |
| Liu LK et al. 2010 | Vimentin (VIM) | (+) | China.<br>Retrospective.<br>IHC. | The high expression of VIM and low expression of CDH1 were associated with survival and were independent prognostic factors in multivariate analyses. |
|  | Cadherin-1 (CDH1) | (-) |  |  |
| Kawaguchi H et al., 2008 | PDPN | (+) | USA.<br>Retrospective.<br>IHC. | Together with histology, may serve as a powerful biomarker to predict the risk for oral cancer development in patients with oral leukoplakia. |
| Pukkila M et al., 2007 | VCAN protein (VCAN) | (+) | Finland.<br>Retrospective.<br>IHC. | Correlated with both increased risk for disease recurrence and shortened survival. High stromal expression may thus be considered an independent and adverse prognostic marker in OSCC. |
| Endo K et al. 2006 | E3 ubiquitin-protein ligase AMFR (AMFR) | (+) | Japan.<br>Retrospective.<br>IHC. | Is valuable in identifying patients at high risk for tongue SCC recurrences |
| **Reprogramming of energy metabolism** |  |  |  |  |
| Hamada T et al., 2012 | Mucin-1 (MUC1) | (+) | Japan.<br>Retrospective.<br>IHC. | Is a risk factor for subsequent lymph node metastasis in patients with OSCC and therefore may represent an indication for elective neck dissection |
| Eckert AW et al. 2011 | Hypoxia-inducible factor 1-α (HIF1A) or HIF-1α and Solute carrier family 2, facilitated glucose transporter member 1 or GLUT-1 (SLC2A1) combined | (+) | Germany.<br>Retrospective.<br>IHC. | Coexpression of high levels of HIF1A and SLC2A1 is significantly correlated with prognosis in OSCC patients. |
| Eckert AW et al., 2010 | HIF1A | (+) | Germany.<br>Retrospective.<br>IHC. | Immunohistochemical detection appears to improve diagnosis and to provide prognostic information in addition to the TNM – system and histological grade of OSCC. |
| Fillies T et al., 2005 | HIF1A | (-) | Germany.<br>Retrospective.<br>IHC. | Overexpression is an indicator of favorable prognosis in T1 and T2 SCC of the oral floor. Node negative patients lacking expression may therefore be considered for adjuvant radiotherapy. |
| **Tumor-promoting inflammation** |  |  |  |  |
| Kwon M et al., 2015 | Interleukin-4 receptor subunit alpha (IL4R) | (+) | Korea. Retrospective. IHC. | High expression of IL4R correlated with increased recurrence, while high IL13RA1 expression had an inverse relationship to recurrence and disease-specific survival in OSCC patients. |
|  | Interleukin-13 receptor subunit alpha-1 (IL13RA1) | (+) |  |  |
| Fujita Y et al. 2014 | Interleukin-8 (CXCL8) | (+) | Japan. Retrospective. IHC. | These factors in addition to N status may have prognostic value in patients with resectable OSCC. |
|  | Scavenger receptor cysteine-rich type 1 protein M130 (CD163) | (+) |  |  |
| Lai WM et al. 2013 | Myeloperoxidase (MPO) | (+) | Taiwan. Retrospective. IHC. | Higher MPO expression in buccal mucosal SCC is a risk factor for second primary tumors. |
| Huang SF et al. 2012 | Serpin B3 (SERPINB3) and C-reactive protein (CRP) combined | (+) | Taiwan. Retrospective. Immunoassay. | High levels of both preoperative SERPINB3 and CRP levels act as a predictor for DFS and OS. |

The articles are grouped according to the hallmarks of cancer. \*UniProt Knowledgebase or common name. HGNC name between parentheses. (+) Up-regulated/overexpressed, (-) Down-regulated/down-expressed, CSS, cause-specific survival; OS, overall survival; DFS, disease free survival; IHC, immunohistochemistry; FISH, fluorescence in situ hybridization.

The included studies were conducted in Poland, India, Germany, Taiwan, Korea, Japan, Australia, Spain, China, Portugal, Brazil, UK, USA and Finland. Variable cohort sizes were used, ranging from 34 to 208 patients. *n*, outcome event number, statistical test, CIs and p-values, risk values and Google scholar citations were extracted (see Supplemental file S1). The main results of the included articles are summarized in Table 2. The biomarker high *vs*. low levels was defined differently in each study.

**Table 2:**
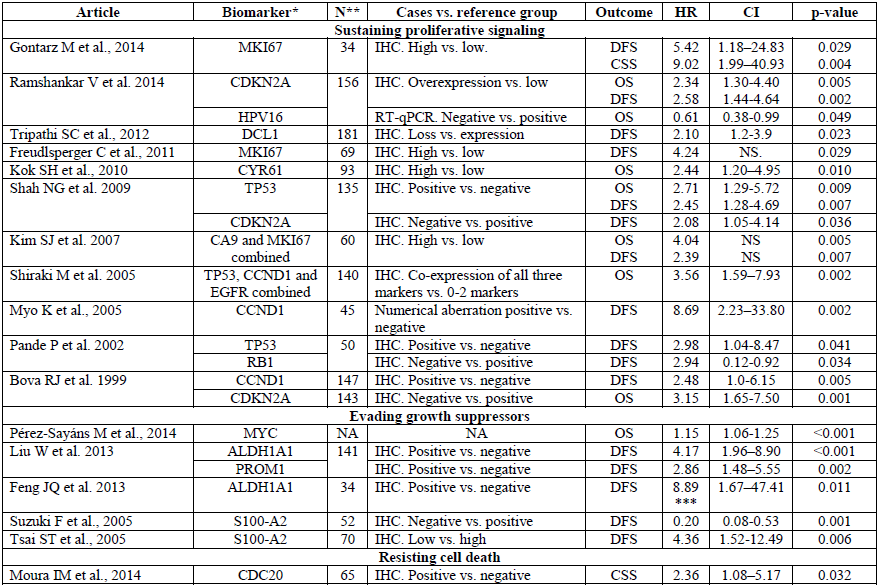
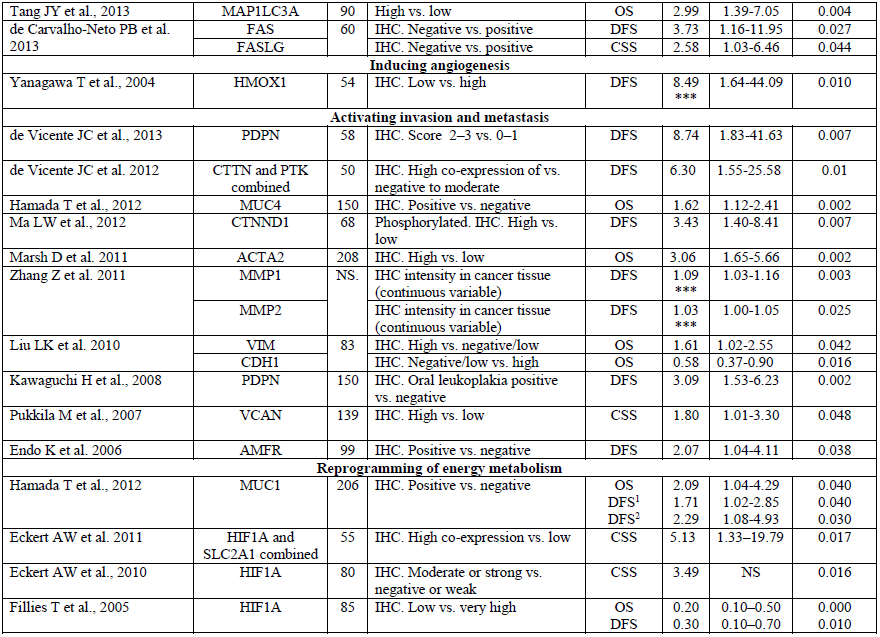
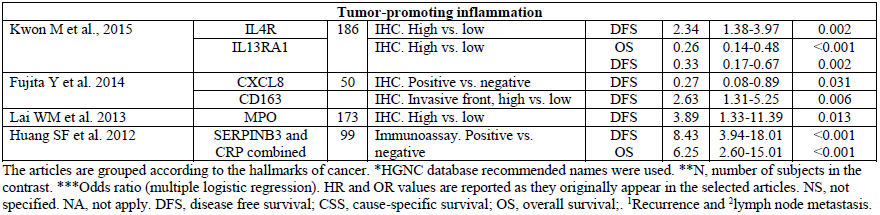
Data extracted from selected studies.

Fourteen clinicopathologic group factors were incorporated in 48 multivariate analyses (38 studies generated 48 significant models and 210 covariables). The most commonly included prognostic factors for model adjustment were the histopathological features (excluding the WHO histological differentiation degree) in 30 models (62,5%), protein (27 models, 53,3%) AJCC clinical stage (22 models, 45,8%) and WHO histological differentiation degree (21 models, 43,8%) (Figure 2). For complete details, see Supplemental file S1.

**Fig. 2.**
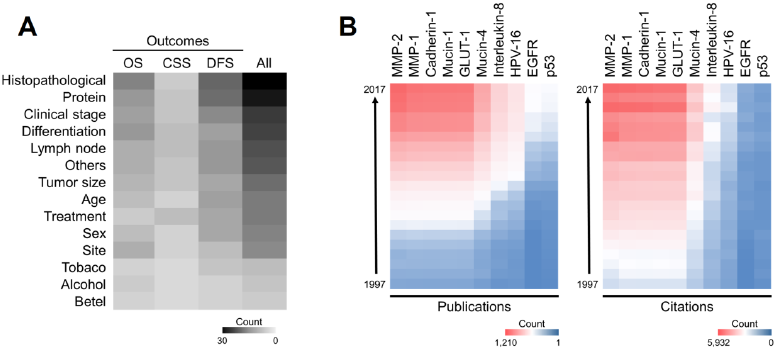
A. Adjustment variables. Frequencies with which adjustments were performed for OSCC outcomes. The heat map combines the most frequent factors for adjustments and survival models. The most commonly included factor was “histopathological features” (excluding the WHO histological differentiation degree). Higher numbers represent intense and saturated colors. B. Trends in oral cancer biomarkers (top ten). Compared with other biomarkers, MMP-2 is the most researched field with 15,057 publications and 46,368 citations (1997-2017). MM-2 is followed by MMP-1 (14,650 publications/43,762 citations) and cadherin-1 (14,531/43,422).

### Quality of study reports: studies do not clear determine the sample size

The result of this agreement was 0.87, which is classified as almost perfect. Differences were resolved by consensus. Most study analyses reported details of the objective/hypothesis, patient source, population characteristics, assay method, cut-off point, and relationship of the potential marker to standard prognostic variables, as well as discussed the implications for future research and clinical value (for details, see Supplemental file S2). Notably, no study referred to a statistical sample size, which is key for biomarker validation.

### Proposed OSCC biomarkers

None of the studied molecules presented an analysis of validation, so we called them “potential biomarkers”. A narrative review of the proposed biomarkers is presented in Table 3.

**Table 3.** Overview of proposed biomarkers

| **Name\*** | **Biological processes** | **Cancer context** |
| --- | --- | --- |
| MKI67 | Cell cycle, cell proliferation. | Marker of the growth fraction for a certain cell population [1]. The labelling index is considered one of the best prognostic factors of the survival rate and recurrence [2]. |
| CDKN2A | Cell cycle, cell cycle arrest. | This gene is frequently mutated or deleted in a wide variety of tumors, and is known to be an important tumor suppressor gene [3]. |
| HPV16 | High-risk HPV type. | Is emerging as an important factor in the rise of oropharyngeal tumors affecting non-smokers in developed countries. Patients with HPV(+) tumors demonstrated favorable outcomes compared to TP53 mutants and 11q13/*CCND1*-amplified tumors [4]. |
| DLC1 | Negative regulation of cell proliferation and migration. | Acts as a tumour suppressor in a number of common cancers, including liver cancer [5]. |
| CYR61 | Regulation of cell growth and adhesion. | Can function as an oncogene or a tumour suppressor, depending on the origin of the cancer [6]. |
| TP53 | Cell cycle, cell cycle arrest. | Tumor-suppressor protein. Mutations in this gene are associated with a variety of human cancers [3]. |
| CA9 | Response to hypoxia. | Is the most widely expressed gene in response to hypoxia. Its role in intracellular pH maintenance represents the means by which cancer cells adapt to the toxic conditions of the extracellular environment [7]. |
| CCND1 | Cell cycle, cell division. | Is frequently deregulated in cancer and is a biomarker of cancer phenotype and disease progression [8]. |
| EGFR | Positive regulation of cell proliferation. | EGFR overexpression is a significant finding in cancer, particularly in head and neck cancer, where it is also associated with a poor prognosis [9]. |
| RB1 | Cell cycle, cell cycle arrest. | Tumor-suppressor protein. Defects in this gene are a cause of childhood cancer retinoblastoma (RB), bladder cancer, and osteogenic sarcoma [3]. |
| MYC | Positive regulation of cell proliferation. | Its oncogenic reputation stems from its frequent deregulation in a host of human cancers and from a suite of activities that place this protein at the nexus of cell growth, proliferation, metabolism, and genome stability [10]. |
| ALDH1A1 | Ethanol oxidation. | Play a key role in the regulation of growth and differentiation of both normal tissue stem cells and cancer stem cells [11]. |
| PROM1 | Retina layer formation. | Maintaining stem cell properties by suppressing differentiation [3]. |
| S100-A2 | Endothelial cell migration. | In epithelial tissue, S100-A2 expression is decreased remarkably in tumours compared with normal specimens [12]. S100-A2 promotes p53 transcriptional activity, and its loss of expression has been associated with a poorer prognosis and shorter survival [13]. |
| CDC20 | Cell cycle, positive regulation of cell proliferation. | The role of CDC20 expression in tumours is not known, but many studies have reported that CDC20 regulates apoptosis, leading to genetically instability [14]. |
| MAP1LC3A | Autophagy. | Strong positive expression in the peripheral area of pancreatic cancer tissue had a shorter overall and disease-free survival; correlations with tumour size, poor differentiation, blood vessel infiltration and tumour necrosis were noted [15]. |
| FAS | Apoptotic process. | Cancer cells can never lose FAS or FASLG. FAS and/or FASLG expression promotes tumor growth and favors the establishment of tumor metastases [16]. |
| FASLG |  |  |
| HMOX1 | Angiogenesis. | Many human tumours produce HMOX1, and its expression is usually higher in cancer cells than in surrounding healthy tissues [17]. |
| PDPN | Lymphangiogenesis. | Is commonly used in the identification of lymphatic endothelial differentiation in vascular endothelial neoplasms and lymphatic invasion by tumours [18]. Recent evidence have identified podoplanin as a marker of cancer-associated fibroblasts [19]. |
| CTTN | Cell motility and focal adhesion assembly. | Is overexpressed in breast cancer and squamous cell carcinomas of the head and neck [3]. |
| PTK2 | Angiogenesis. | Promotes tumor progression and metastasis through effects on cancer cells, as well as stromal cells of the tumor microenvironment [20] |
| MUC4 | Cell adhesion. | An aberrant expression of MUC4 has been reported in various carcinomas [21]. |
| CTNND1 | Cell adhesion. | Evidence is emerging that complete loss, downregulation or mislocalization of CTNND1 correlates with the progression of different types of human tumours [22]. |
| ACTA2 | Mesenchyme migration. | Patients with lung adenocarcinomas and high ACTA2 expression showed significantly enhanced distant metastasis and unfavorable prognosis [23]. |
| MMP1 | Proteolysis. | Imbalance between matrix metalloproteinases and their inhibitors play the important role in progression of head and neck cancer [24]. |
| MMP2 | Angiogenesis, response to hypoxia and proteolysis. |  |
| VIM | Movement of cell or subcellular component. | Has been recognized as a marker for epithelial-mesenchymal transition. Overexpression in cancer correlates well with accelerated tumor growth, invasion, and poor prognosis [25]. |
| CDH1 | Cell adhesion. | Loss of function of this gene is thought to contribute to cancer progression by increasing proliferation, invasion, and/or metastasis [3]. |
| VCAN | Cell adhesion. | Is strongly associated with a poor outcome for many different cancers. Depending on the cancer nature, is expressed either by cancer cells themselves or by stromal cells surrounding the tumour [26]. |
| AMFR | Movement of cell or subcellular component. | Is a tumor motility-stimulating protein secreted by tumor cells [3]. |
| MUC1 | DNA damage response, signal transduction by p53 class mediator resulting in cell cycle arrest | Is aberrantly glycosylated and overexpressed in various epithelial cancers and plays a crucial role in the progression of the disease [27]. MUC1 is often used as a diagnostic marker for metastatic progression [28]. |
| HIF1A | Angiogenesis, response to hypoxia. | Up-regulates the expression of proteins that promote angiogenesis, anaerobic metabolism, and many other survival pathways [29]. |
| SLC2A1 | Glucose transport. | Was significantly correlated with depth of invasion and clinical stage in patients with gastric cancer [30]. |
| IL4R | Immune system process and regulation of cell proliferation | The IL4/IL4R signaling axis is a strong promoter of pro-metastatic phenotypes in epithelial cancer cells including enhanced migration, invasion, survival, and proliferation [31]. |
| IL13RA1 | Cell surface receptor signaling pathway. | Glioblastoma samples presented higher IL13RA1 and IL13RA2 expression levels compared to lower grades astrocytomas and non-neoplastic cases [32]. |
| CXCL8 | Angiogenesis, movement of cell or subcellular component and chemotaxis | Neovascularisation is now recognised as a critical function of CXCL8 in the tumour microenvironment [33]. |
| CD163 | Inflammatory response. | Could be used as a general anti-inflammatory myeloid marker with prognostic impact for breast cancer patients [34]. |
| MPO | Defense response. | Myeloperoxidase-positive cell infiltration in colorectal carcinogenesis is an indicator of colorectal cancer risk [35]. |
| SERPINB3 | Positive regulation of cell proliferation. | Promotes oncogenesis and epithelial-mesenchymal transition [36] |
| CRP | Inflammatory response. | Patients with a high baseline CRP had a greater risk of early death compared with those with low CRP levels [37]. |

\*HGNC database recommended names were used. \*\*Representative processes from QuickGo (http://www.ebi.ac.uk/QuickGO).

### Trends: potential biomarkers with more publications and citations

To explore the publication trends in our OSCC potential protein biomarkers, we searched the scholarly literature in SciCurve Open. SciCurve uses PubMed’s library of 23 million references to generate visually pleasing graphs and curves that help grasp trends in the literature [53]. It is associated with the following main functionalities: publications, citations, most prolific authors and countries.

According to Figure 3, MMP-2 is the most researched field, followed by MMP-1, cadherin-1 and mucin-1. The countries with the largest contributions are the USA, Japan and China.

## DISCUSSION

We have summarized the results on the association between biomarkers and oral cancer outcomes using a systematic review. Overall, our results suggest 41 prognostic molecules involved with OSCC endpoints. These markers may be candidates for long-term studies.

OSCC is the most relevant epithelial malignancy for dental surgeons. It has late clinical detection and poor prognosis, and the available therapeutic alternatives are highly expensive and disfiguring [54].

OSCC is a very complex subtype of cancer with high heterogeneity [55]. Several risk factors are implicated in its aetiology, among which tobacco, alcohol, viruses and diet are highlighted [2]. These factors related to genetic inheritance may have a carcinogenic effect on the normal cells of the respiratory and digestive systems. This type of carcinoma can occur anywhere in the mouth, although the most affected sites are the tongue, lower lip and mouth floor [2, 56]. These regions are great facilitators of carcinoma spreading to regional lymph nodes and/or distant organs [57]. At present, the diagnosis of OSCC is based on comprehensive clinical examination and histological analysis of suspicious areas [58]. Recently, The Cancer Genome Atlas (TCGA) showed that a large dataset of proteomics/genomics did not improve the prognosis potential of classic clinical variables in patients with different types of cancer [59]. Some studies seeking biomarkers in oral cancer are still in the discovery phase, requiring validation to be accepted in clinical practice.

Currently, biomarkers are a subject of particular interest because they may represent the most important part in the diagnosis step. In the future, specific and personalised diagnostics can guide treatment against the disease and consequently improve the chance of curing the disease.

In response to the need for tumor biomarkers for OSCC that can be readily evaluated in routine clinical practice, we performed a systematic review (PubMed keyword*-*base query) of the published literature to identify single or multiple biomarkers for OSCC outcomes: overall survival, disease-free survival, relapse-free survival and cause-specific survival. The main finding was the identification of 38 studies describing multivariate survival analysis for 41 biomarkers. From these articles, MMP-2, MMP-1, cadherin-1, mucin-1, GLUT-1 (SLC2A1), mucin-4, interleukin-8, HPV-16, EGFR and p53 have received great interest from the scientific community. Of these, up to now, it is accepted that the HPV status have a clinical utility [60], suggesting that HPV positive head and neck squamous cell carcinomas form a distinct clinical entity with better treatment outcome [61].

The malignant progression to OSCC is characterized by the acquisition of progressive and uncontrolled growth of tumor cells. Predicting whether premalignant lesions will progress to cancer is crucial to make appropriate treatment decisions. The first detectable clinical changes that can indicate that an epithelium is on the way to establish OSCC is the occurrence of malignant disorders, including leukoplakia (most common) [2]. In this context, we emphasize the results associated with Rho GTPase-activating protein 7, retinal dehydrogenase 1/prominin-1 (combined biomarkers), podoplanin, cortactin/focal adhesion kinase 1 (combined biomarkers) and catenin delta-1. These proteins show a potential role as a marker of oral cancer risk and malignant transformation [17, 26-28, 39, 40, 42].

There are thousands of papers reporting cancer biomarker discovery, but only few clinically useful biomarkers have been successfully validated for routine clinical practice [62]. Quality assessment tools have been developed for prognostic studies to help identify study biases and causes of heterogeneity when performing meta-analysis. We chose to use the REMARK reporting guidelines, which provide a useful start for assessing tumor prognostic biomarkers (all included studies were prognostic). We found that the investigations reported an average of 19 of 20 REMARK items. However, all studies failed to report the sample size calculation. In the absence of this calculation, the findings of each research should be interpreted with caution [63]. The sample size requirements that allow the identification of a benefit beyond existing biomarkers are even more demanding [64].

In our review, none of the articles that created prediction models had internal or external validation. In general, studies recruited cases of OSCC from a clinical setting as well as controls without a clearly defined diagnosis. Under this circumstance, any differences in the biomarker levels between OSCC patients and controls could simply reflect individual differences rather than cancer-related differences. The lack of biomarker validation strategies and standard operating procedures for sample selection in the included studies represent an important pitfalls and limitations, leading us to use the term "potential biomarkers" instead of biomarker in our article title.

It is important to highlight that our research searched only one database, which means that only studies available in MEDLINE were included. Additionally, due to the heterogeneity among the studies, a meta-analysis that combined the results of different studies could not be performed. In addition, our research included results from observational studies, and their evaluation may have been problematic if the confounder variables were not adjusted because they were not measured [65].

## CONCLUSION

Recent research in OSCC has identified a multitude of potential markers that have a significant role in prognosis. In this systematic review, despite the inherent limitations, we identified several potential biomarkers of particular interest that appear to carry prognostic significance. Considering the validation step as a process of assessing the biomarker and its measurement performance characteristics, and determine the range of conditions under which this biomarker can provide reproducible data [9], our results show biomarkers in the discovery phase, thereby leading us to call them OSCC “potential biomarkers”. Nevertheless, it is urgent to apply validation methods to provide clinically useful oral cancer biomarkers.

## ACKNOWLEDGMENT

CONICYT Becas-Chile Scholarship 8540/2014, PNPD/CAPES 33003033009P4, and FAPESP Grants 2016/07846-0, 2014/06485-9 and 2015/12431-1, supported this work.

## REFERENCES

[1] Chi AC, Day TA, Neville BW. Oral cavity and oropharyngeal squamous cell carcinoma--an update. CA Cancer J Clin. 2015;65:401–21.

[2] Rivera C. Essentials of oral cancer. Int J Clin Exp Pathol. 2015;8:11884–94.

[3] Huss R. Tissue-based biomarkers. In: Günther C HA, Huss R, editor. Advances in Pharmaceutical Cell Therapy: Principles of Cell-Based Biopharmaceuticals. Singapore: World Scientific; 2015. p. 408.

[4] Henry NL, Hayes DF. Cancer biomarkers. Mol Oncol. 2012;6:140–6.

[5] Mishra A, Verma M. Cancer biomarkers: are we ready for the prime time? Cancers (Basel). 2010;2:190–208.

[6] Ballman KV. Biomarker: Predictive or Prognostic? J Clin Oncol. 2015;33:3968–71.

[7] Buyse M, Michiels S, Sargent DJ, lGrothey A, Matheson A, de Gramont A. Integrating biomarkers in clinical trials. Expert Rev Mol Diagn. 2011;11:171–82.

[8] Gosho M, Nagashima K, Sato Y. Study Designs and Statistical Analyses for Biomarker Research. Sensors (Basel, lSwitzerland). 2012;12:8966–86.

[9] Hunter DJ, Losina E, Guermazi A, lBurstein D, Lassere MN, Kraus V. A pathway and approach to biomarker validation and qualification for osteoarthritis clinical trials. Curr Drug Targets. 2010;11:536–45.

[10] Haddaway N. The importance of meta-analysis and systematic review: How research legacy can be maximized through adequate reporting. LSE Impact of Social Sciences; 2015.

[11] Mallett S, Timmer A,Sauerbrei W, lAltman DG. Reporting of prognostic studies of tumor markers: a review of published articles in relation to REMARK guidelines. Br J Cancer. 2010;102:173–80.

[12] Altman DG, McShane LM, Sauerbrei W, lTaube SE. Reporting Recommendations for Tumor Marker Prognostic Studies (REMARK): explanation and elaboration. PLoS Med. 2012;9:e1001216.

[13] Shiba B, Nabakumar B. Research2. 0: Recent Tools and Technology that’s Makes Web as a Research Platform. Global Journal of Multidisciplinary Studies. 2015;4.

[14] Hanahan D, Weinberg RA. Hallmarks of cancer: the next generation. Cell. 2011;144:646– 74.

[15] Gontarz M, Wyszynska-Pawelec G, Zapala J, lCzopek J, Lazar A, Tomaszewska R. Proliferative index activity in oral squamous cell carcinoma: indication for postoperative radiotherapy? Int J Oral Maxillofac Surg. 2014;43:1189–94.

[16] Freudlsperger C, Rohleder SE, Reinert S, lHoffmann J. Predictive value of high Ki-67 expression in stage I oral squamous cell carcinoma specimens after primary surgery. Head Neck. 2011;33:668–72.

[17] Tripathi SC, Kaur J, Matta A, lGao X, Sun B, Chauhan SS, et al. Loss of DLC1 is an independent prognostic factor in patients with oral squamous cell carcinoma. Mod Pathol. 2012;25:14–25.

[18] Kok SH, Chang HH, Tsai JY, lHung HC, Lin CY, Chiang CP, et al. Expression of Cyr61 (CCN1) in human oral squamous cell carcinoma: An independent marker for poor prognosis. Head Neck. 2010;32:1665–73.

[19] Perez-Sayans M, Suarez-Penaranda JM, Padin-Iruegas E, lGayoso-Diz P, Reis-De Almeida M, Barros-Angueira F, et al. Quantitative determination of c-myc facilitates the assessment of prognosis of OSCC patients. Oncol Rep. 2014;31:1677-82.

[20] Suzuki F, Oridate N, Homma A, lNakamaru Y, Nagahashi T, Yagi K, et al. S100A2 expression as a predictive marker for late cervical metastasis in stage I and II invasive squamous cell carcinoma of the oral cavity. Oncol Rep. 2005;14:1493–8.

[21] Tsai ST, Jin YT, Tsai WC, lWang ST, Lin YC, Chang MT, et al. S100A2, a potential marker for early recurrence in early-stage oral cancer. Oral Oncol. 2005;41:349–57.

[22] Moura IM, Delgado ML, Silva PM, lLopes CA, do Amaral JB, Monteiro LS, et al. High CDC20 expression is associated with poor prognosis in oral squamous cell carcinoma. J Oral Pathol Med. 2014;43:225–31.

[23] Tang JY, Hsi E, Huang YC, lHsu NC, Chu PY, Chai CY. High LC3 expression correlates with poor survival in patients with oral squamous cell carcinoma. Hum Pathol. 2013;44:2558–62.

[24] Yanagawa T, Omura K, Harada H, lNakaso K, Iwasa S, Koyama Y, et al. Heme oxygenase-1 expression predicts cervical lymph node metastasis of tongue squamous cell carcinomas. Oral Oncol. 2004;40:21–7.

[25] Hamada T, Wakamatsu T, Miyahara M, lNagata S, Nomura M, Kamikawa Y, et al. MUC4: a novel prognostic factor of oral squamous cell carcinoma. Int J Cancer. 2012;130:1768–76.

[26] de Vicente JC, Rodrigo JP, Rodriguez-Santamarta T, lLequerica-Fernandez P, Allonca E, Garcia-Pedrero JM. Podoplanin expression in oral leukoplakia: tumorigenic role. Oral Oncol. 2013;49:598–603.

[27] Kawaguchi H, El-Naggar AK, Papadimitrakopoulou V, lRen H, Fan YH, Feng L, et al. Podoplanin: a novel marker for oral cancer risk in patients with oral premalignancy. J Clin Oncol. 2008;26:354–60.

[28] Ma LW, Zhou ZT, He QB, lJiang WW. Phosphorylated p120-catenin expression has predictive value for oral cancer progression. J Clin Pathol. 2012;65:315–9.

[29] Pukkila M, Kosunen A, Ropponen K, lVirtaniemi J, Kellokoski J, Kumpulainen E, et al. High stromal versican expression predicts unfavourable outcome in oral squamous cell carcinoma. J Clin Pathol. 2007;60:267–72.

[30] Hamada T, Nomura M, Kamikawa Y, lYamada N, Batra SK, Yonezawa S, et al. DF3 epitope expression on MUC1 mucin is associated with tumor aggressiveness, subsequent lymph node metastasis, and poor prognosis in patients with oral squamous cell carcinoma. Cancer. 2012;118:5251–64.

[31] Eckert AW, Schutze A, Lautner MH, lTaubert H, Schubert J, Bilkenroth U. HIF-1alpha is a prognostic marker in oral squamous cell carcinomas. Int J Biol Markers. 2010;25:87–92.

[32] Fillies T, Werkmeister R, van Diest PJ, Brandt B, Joos U, Buerger H. HIF1-alpha overexpression indicates a good prognosis in early stage squamous cell carcinomas of the oral floor. BMC Cancer. 2005;5:84.

[33] Myo K, Uzawa N, Miyamoto R, Sonoda I, Yuki Y, Amagasa T. Cyclin D1 gene numerical aberration is a predictive marker for occult cervical lymph node metastasis in TNM Stage I and II squamous cell carcinoma of the oral cavity. Cancer. 2005;104:2709–16.

[34] Kwon M, Kim JW, Roh JL, Park Y, Cho KJ, Choi SH, et al. Recurrence and cancer-specific survival according to the expression of IL-4Ralpha and IL-13Ralpha1 in patients with oral cavity cancer. Eur J Cancer. 2015;51:177–85.

[35] Fujita Y, Okamoto M, Goda H, Tano T, Nakashiro K, Sugita A, et al. Prognostic significance of interleukin-8 and CD163-positive cell-infiltration in tumor tissues in patients with oral squamous cell carcinoma. PLoS One. 2014;9:e110378.

[36] Ramshankar V, Soundara VT, Shyamsundar V, Ramani P, Krishnamurthy A. Risk stratification of early stage oral tongue cancers based on HPV status and p16 immunoexpression. Asian Pac J Cancer Prev. 2014;15:8351–9.

[37] Lai WM, Chen CC, Lee JH, Chen CJ, Wang JS, Hou YY, et al. Second primary tumors and myeloperoxidase expression in buccal mucosal squamous cell carcinoma. Oral Surg Oral Med Oral Pathol Oral Radiol. 2013;116:464–73.

[38] de Carvalho-Neto PB, dos Santos M, de Carvalho MB, Mercante AM, dos Santos VP, Severino P, et al. FAS/FASL expression profile as a prognostic marker in squamous cell carcinoma of the oral cavity. PLoS One. 2013;8:e69024.

[39] Liu W, Wu L, Shen XM, Shi LJ, Zhang CP, Xu LQ, et al. Expression patterns of cancer stem cell markers ALDH1 and CD133 correlate with a high risk of malignant transformation of oral leukoplakia. Int J Cancer. 2013;132:868–74.

[40] Feng JQ, Xu ZY, Shi LJ, Wu L, Liu W, Zhou ZT. Expression of cancer stem cell markers ALDH1 and Bmi1 in oral erythroplakia and the risk of oral cancer. J Oral Pathol Med. 2013;42:148–53.

[41] Huang SF, Wei FC, Liao CT, Wang HM, Lin CY, Lo S, et al. Risk stratification in oral cavity squamous cell carcinoma by preoperative CRP and SCC antigen levels. Ann Surg Oncol. 2012;19:3856–64.

[42] de Vicente JC, Rodrigo JP, Rodriguez-Santamarta T, Lequerica-Fernandez P, Allonca E, Garcia-Pedrero JM. Cortactin and focal adhesion kinase as predictors of cancer risk in patients with premalignant oral epithelial lesions. Oral Oncol. 2012;48:641–6.

[43] Eckert AW, Lautner MH, Schutze A, Taubert H, Schubert J, Bilkenroth U. Coexpression of hypoxia-inducible factor-1alpha and glucose transporter-1 is associated with poor prognosis in oral squamous cell carcinoma patients. Histopathology. 2011;58:1136–47.

[44] Marsh D, Suchak K, Moutasim KA, Vallath S, Hopper C, Jerjes W, et al. Stromal features are predictive of disease mortality in oral cancer patients. J Pathol. 2011;223:470–81.

[45] Zhang Z, Pan J, Li L, Wang Z, Xiao W, Li N. Survey of risk factors contributed to lymphatic metastasis in patients with oral tongue cancer by immunohistochemistry. J Oral Pathol Med. 2011;40:127–34.

[46] Liu LK, Jiang XY, Zhou XX, Wang DM, Song XL, Jiang HB. Upregulation of vimentin and aberrant expression of E-cadherin/beta-catenin complex in oral squamous cell carcinomas: correlation with the clinicopathological features and patient outcome. Mod Pathol. 2010;23:213– 24.

[47] Shah NG, Trivedi TI, Tankshali RA, Goswami JV, Jetly DH, Shukla SN, et al. Prognostic significance of molecular markers in oral squamous cell carcinoma: a multivariate analysis. Head Neck. 2009;31:1544–56.

[48] Kim SJ, Shin HJ, Jung KY, Baek SK, Shin BK, Choi J, et al. Prognostic value of carbonic anhydrase IX and Ki-67 expression in squamous cell carcinoma of the tongue. Jpn J Clin Oncol. 2007;37:812–9.

[49] Endo K, Shirai A, Furukawa M, Yoshizaki T. Prognostic value of cell motility activation factors in patients with tongue squamous cell carcinoma. Hum Pathol. 2006;37:1111–6.

[50] Shiraki M, Odajima T, Ikeda T, Sasaki A, Satoh M, Yamaguchi A, et al. Combined expression of p53, cyclin D1 and epidermal growth factor receptor improves estimation of prognosis in curatively resected oral cancer.Mod Pathol. 2005;18:1482–9.

[51] Pande P, Soni S, Kaur J, Agarwal S, Mathur M, Shukla NK, et al. Prognostic factors in betel and tobacco related oral cancer. Oral Oncol. 2002;38:491–9.

[52] Bova RJ, Quinn DI, Nankervis JS, Cole IE, Sheridan BF, Jensen MJ, et al. Cyclin D1 and p16INK4A expression predict reduced survival in carcinoma of the anterior tongue. Clin Cancer Res. 1999;5:2810–9.

[53] Connected Researchers. SciCurve: revealing life science’s curves. 2014.

[54] Rivera C. Opportunities for biomarkers with potential clinical use in oral cancer. Medwave. 2015;15:e6186.

[55] Bavle RM, Venugopal R, Konda P, Muniswamappa S, Makarla S. Molecular Classification of Oral Squamous Cell Carcinoma. J Clin Diagn Res. 2016;10:ZE18–ZE21.

[56] Rivera C, Venegas B. Histological and molecular aspects of oral squamous cell carcinoma (Review). Oncol Lett. 2014;8:7–11.

[57] Donaduzzi LC, De-Conto F, Kuze LS, Rovani G, Flores ME, Pasqualotti A. Occurrence of contralateral lymph neck node metastasis in patients with squamous cell carcinoma of the oral cavity. J Clin Exp Dent. 2014;6:e209–13.

[58] Bano S, David MP, Indira A. Salivary Biomarkers for Oral Squamous Cell Carcinoma: An Overview. IJSS. 2015;1:39.

[59] Yuan Y, Van Allen EM, Omberg L, Wagle N, Amin-Mansour A, Sokolov A, et al. Assessing the clinical utility of cancer genomic and proteomic data across tumor types. 2014;32:644–52.

[60] The Cancer Genome Atlas Network. Comprehensive genomic characterization of head and neck squamous cell carcinomas. Nature. 2015;517:576–82.

[61] Dok R, Nuyts S. HPV Positive Head and Neck Cancers: Molecular Pathogenesis and Evolving Treatment Strategies. Cancers (Basel). 2016;8.

[62] Poste G.Bring on the biomarkers. Nature. 2011;469:156–7.

[63] Faber J, Fonseca LM. How sample size influences research outcomes. Dental Press J Orthod. 2014;19:27–9.

[64] Levy MM. Biomarkers in the Critically Ill Patient, an Issue of Critical Care Clinics: Elsevier-Health Sciences Division; 2011.

[65] Shrier I, Boivin JF, Steele RJ, Platt RW, Furlan A, Kakuma R, et al. Should meta-analyses of interventions include observational studies in addition to randomized controlled trials? A critical examination of underlying principles. Am J Epidemiol. 2007;166:1203–9.

## TABLE 3 REFERENCES

[1] Scholzen T, Gerdes J. The Ki-67 protein: from the known and the unknown. J Cell Physiol. 2000;182:311–22.

[2] Rivera C, Venegas B. Histological and molecular aspects of oral squamous cell carcinoma (Review). Oncol Lett. 2014;8:7–11.

[3] Uhlen M, Fagerberg L, Hallstrom BM, Lindskog C, Oksvold P, Mardinoglu A, et al. Proteomics. Tissue-based map of the human proteome. Science. 2015;347:1260419.

[4] The Cancer Genome Atlas Network. Comprehensive genomic characterization of head and neck squamous cell carcinomas. Nature. 2015;517:576–82.

[5] Lukasik D, Wilczek E, Wasiutynski A, Gornicka B. Deleted in liver cancer protein family in human malignancies (Review). Oncol Lett. 2011;2:763–8.

[6] Jeong D, Heo S, Sung Ahn T, Lee S, Park S, Kim H, et al. Cyr61 expression is associated with prognosis in patients with colorectal cancer. BMC Cancer. 2014;14:164.

[7] Benej M, Pastorekova S, Pastorek J. Carbonic anhydrase IX: regulation and role in cancer. Subcell Biochem. 2014;75:199–219.

[8] Musgrove EA, Caldon CE, Barraclough J, Stone A, Sutherland RL. Cyclin D as a therapeutic target in cancer. Nat Rev Cancer. 2011;11:558– 72.

[9] Oliveira-Silva RJ, Carolina de Carvalho A, de Souza Viana L, Carvalho AL, Reis RM. Anti-EGFR Therapy: Strategies in Head and Neck Squamous Cell Carcinoma. Recent Pat Anticancer Drug Discov. 2016;11:170–83.

[10] Tansey WP. Mammalian MYC proteins and cancer. New Journal of Science. 2014;2014.

[11] Tomita H, Tanaka K, Tanaka T, Hara A. Aldehyde dehydrogenase 1A1 in stem cells and cancer. Oncotarget. 2016;7:11018–32.

[12] Nagy N, Brenner C, Markadieu N, Chaboteaux C, Camby I, Schafer BW, et al. S100A2, a putative tumor suppressor gene, regulates in vitro squamous cell carcinoma migration. Lab Invest. 2001;81:599–612.

[13] Salama I, Malone PS, Mihaimeed F, Jones JL. A review of the S100 proteins in cancer. Eur J Surg Oncol. 2008;34:357–64.

[14] Wan L, Tan M, Yang J, Inuzuka H, Dai X, Wu T, et al. APC(Cdc20) suppresses apoptosis through targeting Bim for ubiquitination and destruction. Dev Cell. 2014;29:377–91.

[15] Fujii S, Mitsunaga S, Yamazaki M, Hasebe T, Ishii G, Kojima M, et al. Autophagy is activated in pancreatic cancer cells and correlates with poor patient outcome. Cancer Sci. 2008;99:1813–9.

[16] Peter ME, Hadji A, Murmann AE, Brockway S, Putzbach W, Pattanayak A, et al. The role of CD95 and CD95 ligand in cancer. Cell Death Differ. 2015;22:885–6.

[17] Zhao H, Ozen M, Wong RJ, Stevenson DK. Heme oxygenase-1 in pregnancy and cancer: similarities in cellular invasion, cytoprotection, angiogenesis, and immunomodulation. Front Pharmacol. 2014;5:295.

[18] Ordonez NG. Value of podoplanin as an immunohistochemical marker in tumor diagnosis: a review and update. Appl Immunohistochem Mol Morphol. 2014;22:331–47.

[19] Pula B, Witkiewicz W, Dziegiel P, Podhorska-Okolow M. Significance of podoplanin expression in cancer-associated fibroblasts: a comprehensive review. Int J Oncol. 2013;42:1849–57.

[20] Sulzmaier FJ, Jean C, Schlaepfer DD. FAK in cancer: mechanistic findings and clinical applications. Nat Rev Cancer. 2014;14:598–610.

[21] Chaturvedi P, Singh AP, Batra SK. Structure, evolution, and biology of the MUC4 mucin. Faseb j. 2008;22:966–81.

[22] van Hengel J, van Roy F. Diverse functions of p120ctn in tumors. Biochim Biophys Acta. 2007;1773:78–88.

[23] Lee HW, Park YM, Lee SJ, Cho HJ, Kim DH, Lee JI, et al. Alpha-smooth muscle actin (ACTA2) is required for metastatic potential of human lung adenocarcinoma. Clin Cancer Res. 2013;19:5879–89.

[24] Pietruszewska W, Bojanowska-Pozniak K, Kobos J. Matrix metalloproteinases MMP1, MMP2, MMP9 and their tissue inhibitors TIMP1, TIMP2, TIMP3 in head and neck cancer: an immunohistochemical study. Otolaryngol Pol. 2016;70:32–43.

[25] Satelli A, Li S. Vimentin in cancer and its potential as a molecular target for cancer therapy. Cell Mol Life Sci. 2011;68:3033–46.

[26] Ricciardelli C, Sakko AJ, Ween MP, Russell DL, Horsfall DJ. The biological role and regulation of versican levels in cancer. Cancer Metastasis Rev. 2009;28:233–45.

[27] Nath S, Mukherjee P. MUC1: a multifaceted oncoprotein with a key role in cancer progression. Trends Mol Med. 2014;20:332–42.

[28] Horm TM, Schroeder JA. MUC1 and metastatic cancer: expression, function and therapeutic targeting. Cell Adh Migr. 2013;7:187–98.

[29] Koh MY, Spivak-Kroizman TR, Powis G. HIF-1alpha and cancer therapy. Recent Results Cancer Res. 2010;180:15–34.

[30] Yan S, Wang Y, Chen M, Li G, Fan J. Deregulated SLC2A1 Promotes Tumor Cell Proliferation and Metastasis in Gastric Cancer. Int J Mol Sci. 2015;16:16144–57.

[31] Bankaitis KV, Fingleton B. Targeting IL4/IL4R for the treatment of epithelial cancer metastasis. Clin Exp Metastasis. 2015;32:847–56.

[32] Moretti IF, Silva R, Oba-Shinjo SM, Carvalho POd, Cardoso LC, Castro Id, et al. The impact of interleukin-13 receptor expressions in cell migration of astrocytomas. MedicalExpress. 2015;2.

[33] Liu Q, Li A, Tian Y, Wu JD, Liu Y, Li T, et al. The CXCL8-CXCR1/2 pathways in cancer. Cytokine Growth Factor Rev. 2016;31:61–71.

[34] Medrek C, Ponten F, Jirstrom K, Leandersson K. The presence of tumor associated macrophages in tumor stroma as a prognostic marker for breast cancer patients. BMC Cancer. 2012;12:306.

[35] Roncucci L, Mora E, Mariani F, Bursi S, Pezzi A, Rossi G, et al. Myeloperoxidase-positive cell infiltration in colorectal carcinogenesis as indicator of colorectal cancer risk. Cancer Epidemiol Biomarkers Prev. 2008;17:2291–7.

[36] Sheshadri N, Catanzaro JM, Bott AJ, Sun Y, Ullman E, Chen EI, et al. SCCA1/SERPINB3 promotes oncogenesis and epithelial-mesenchymal transition via the unfolded protein response and IL6 signaling. Cancer Res. 2014;74:6318–29.

[37] Allin KH, Nordestgaard BG. Elevated C-reactive protein in the diagnosis, prognosis, and cause of cancer. Crit Rev Clin Lab Sci. 2011;48:155–70.

